# Deep-RBPPred: Predicting RNA binding proteins in the proteome scale based on deep learning

**DOI:** 10.1101/210153

**Authors:** Jinfang Zheng, Xiaoli Zhang, Xunyi Zhao, Xiaoxue Tong, Xu Hong, Juan Xie, Shiyong Liu

## Abstract

RNA binding protein (RBP) plays an important role in cell processes. Identifying RBPs by computation and experiment are both essential. Recently, RBPPred is proposed in our group to predict RBP with a high performance. However, RBPPred is too slow for that it will generate PSSM matrix as its feature. Herein, we develop a deep learning model called Deep-RBPPred. The model has three advantages comparing to previous models. 1. Deep-RBPPred only needs few physicochemical properties. 2. Deep-RBPPred runs much faster. 3. Deep-RBPPred has a good generalization ability. In the meantime, the performance is still as good as the stats-of-the-art method. In the testing in A. thaliana, S. cerevisiae and H. sapiens proteomics, MCC (AUC) are 0.6077 (0.9421), 0.573 (0.9034) and 0.8141(0.9515) respectively when the score cutoff is set to 0.5. In the verifying in Gerstberger-1538, the SN of our model is 90.38%. The running times are 9s, 7s, 8s and 10s, respectively, for H.sapiens, A.thaliana, S.cerevisiae and Gerstberger-1538 when it is tested in GPU. Deep-RBPPred forecasts 94.65% of 299 new RBP and about 8% higher sensitivity than RBPPred. We also apply deep-RBPPred in 19 eukaryotes proteomics and 11 bacteria proteomics downloaded from Uniprot. The result shows that rate of RBPs in eukaryotes proteome are much higher than bacteria proteome. Testing in 6 proteomics shows the many RBPs may be still undiscovered so far.

## Introduction

RNA binding proteins (RBPs) play important functions in many cellar processes, such as post-transcriptional gene regulation, RNA subcellular localization and alternative splicing. With significant function in biology, many high-throughput experimental techniques have been developed to identify new RBPs in human, mouse and S.cerevisiae (Baltz et al. 2012; Castello et al. 2012; Kwon et al. 2013; Mitchell et al. 2013). Also, many computational methods have been proposed to predict RBPs (Zhao et al. 2011; Yang et al. 2012; Paz et al. 2016; Sharan et al. 2017). Previous computational methods only considered only part features or known RNA binding domain (RBD) which play a significant role in RBP predicting. Based on this consideration, we proposed RBPPred (Zhang and Liu 2017) to address this problem. Benchmarking on datasets shows that RBPPred is better than other approaches (Zhang and Liu 2017). But RBPPred runs much slow because it is required to run blast against a huge protein NR database to generate PSSM matrix. To overcome this shortcoming, we present Deep-RBPPred which is based on deep learning.

In recently years, deep learning technology is used in many aspects in bioinformatics and proved as a power tool (Alipanahi et al. 2015; Kelley et al. 2016; Zeng et al. 2016). In Deep-RBPPred, we apply a deep convolutional neural network to train the RBP predictor instead of SVM. Like RBPPred, we only employ physical-chemical features including hydrophobicity, polarity, normalized van der Waals volume, polarizability, predicted solvent accessibility, side chain’s charge and polarity. These features are used to train the weights of 11 layers convolutional neural network with Tensorflow (Abadi et al. 2016). Finally, the trained model was tested on the independent datasets and predicted RBPs on 30 proteomics.

## Method

### Training set and testing set

In order to train our deep learning model and test the prediction ability, we used two datasets which published in the PRBPred (Zhang and Liu 2017). The training set is consisted of 2780 RBPs and 7093 non-RBPs derived from PDB. And the testing set includes three species: Homo sapiens (H.sapiens), Saccharomyces cerevisiae (S.cerevisiae) and Arabidopsis thaliana (A.thaliana). The positive samples were collected from the Uniprot database, which is retrieval with GO term “RNA binding” to search this protein database and the redundancy sequences are removed by PISCES (Wang and Dunbrack 2003). The negative samples are collected from PDB. The testing dataset contains 967 RBPs and 597 non-RBPs for H. sapiens, 354 RBPs and 135 non-RBPs for S. cerevisiae and 456 RBPs and 37 non-RBPs for A. thaliana, respectively.

### Gerstberger-1538

Gerstberger-1538 is a positive dataset only including human RBPs. This dataset contains 916 experimental identified sequences and 622 computational identified sequences (Gerstberger et al. 2014). We receive the dataset RBPPred (Zhang and Liu 2017).

### Protein features and encoding

The protein was encoded by the approach described in RBPPred (Zhang and Liu 2017). But the evolutionary information and predicted secondary structure are discarded due to the computational time. At last, total of a 160 dimensional vector is encoded to represent each protein sequence including the properties of hydrophobicity, normalized van der Waals volume, polarity and polarizability, solvent accessibility, charge and polarity of side chain.

### Performance evaluation

The performance was evaluated by sensitivity (SN), specificity (SP), precision (PRE), accuracy (ACC), F-measure and Matthews correlation coefficient (MCC) which are defined as follow:

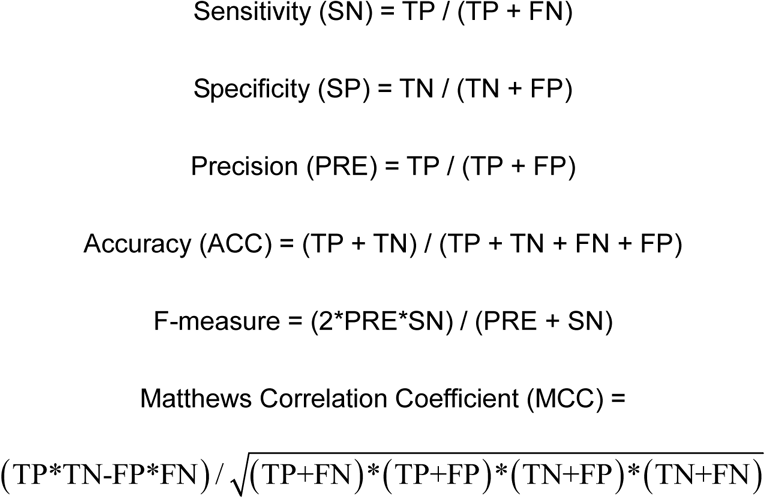

where, TP is true positive, FN is false negative, TN refers to true negative and FP refers to false positive. The receiver operation characteristic (ROC) curve and AUC values are also used to evaluate the performance of model.

### Network structure of Deep-RBPPred

Deep-RBPPred is based on Convolutional Neural Network (CNN) from tensorflow. In Figure 1, it shows the network structure of Deep-RBPPred. The input layer is a size 8×20 feature tensor encoded by proteins. The next layer is a convolution layer with kernel size 2×5, in this layer 32 convolution kernels are set to filter the input features. The third layer is a max pooling layer with size 2×2, the feature size will be reduced to 4×10 after the layer. And the next is a local response normalized layer. This layer is set to increase the generalization ability. The following three layers respectively are convolution layer, max pooling layer and local response normalized layer. Then the feature tensor is flatted to a 640 dimensional vector. The following two layers are fully connections layers with 512 and 256 neurons. The 10^th^ layer is a dropout layer which randomly discards some neurons in the training phase. The final layer is the Softmax layer which is used to classify RNA binding protein or not. The output of this model is a probability score which describes the probability of an RBP. All the activation functions in neurons are ReLU. All the weights in neurons are added a L2 regularization operation. The L2 regularization losses are added to the final loss function. This system is trained to minimize final loss function which is consisted of cross-entropy between the label and probability score and L2 regularization loss of neurons.

**Figure 1.**
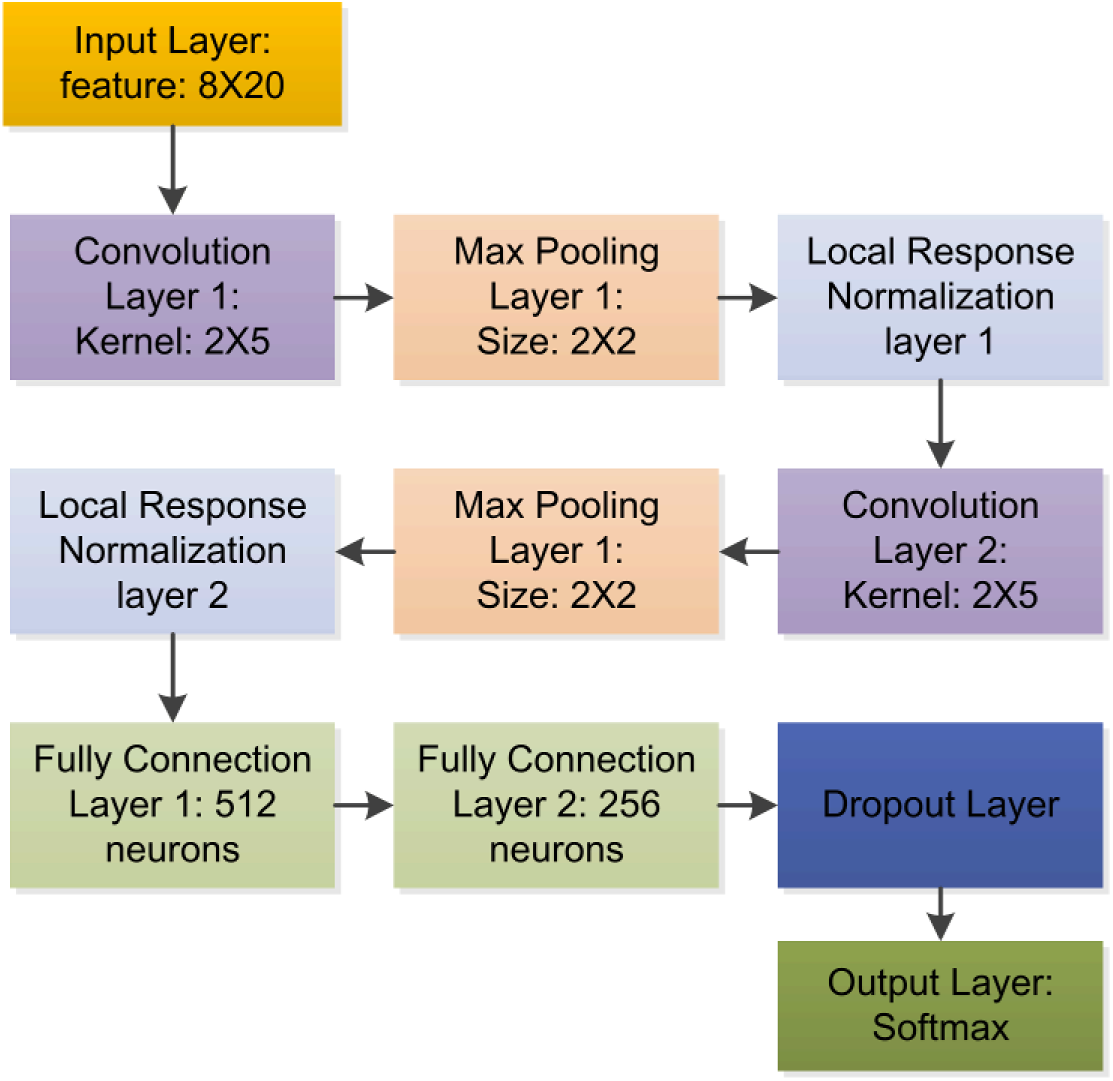
Network structure of Deep-RBPPred. Deep-RBPPred is a CNN network including 11 layers. Convolution Layer and Max Pooling Layer are designed to automatically extract the feature. Local Response Normalization layer and Dropout Layer are designed to avoid over-fitting. Softmax layer is used to classify the protein with probability score.

## Result

### Model training

We total train the model in training sets with 42,000 steps. In each step of model training, 9873 samples are randomly divided into three parts. For training part, the number of sequence is 200. For validating part, the number of sequence is 600. The probability score is chosen as 0.5 to classify the RBP and non-RBP. This model is trained with the ADAM optimizer and stochastic gradient descent. In Figure 2, it shows all results of training steps. The training ACC and validating ACC is increased with the number of training steps. At the last phase, the training ACC and validating ACC is closed to 1.0 and waved around 1.0 with the training steps. This phenomenon indicates that the training model have reached the highest ACC.

**Figure 2.**
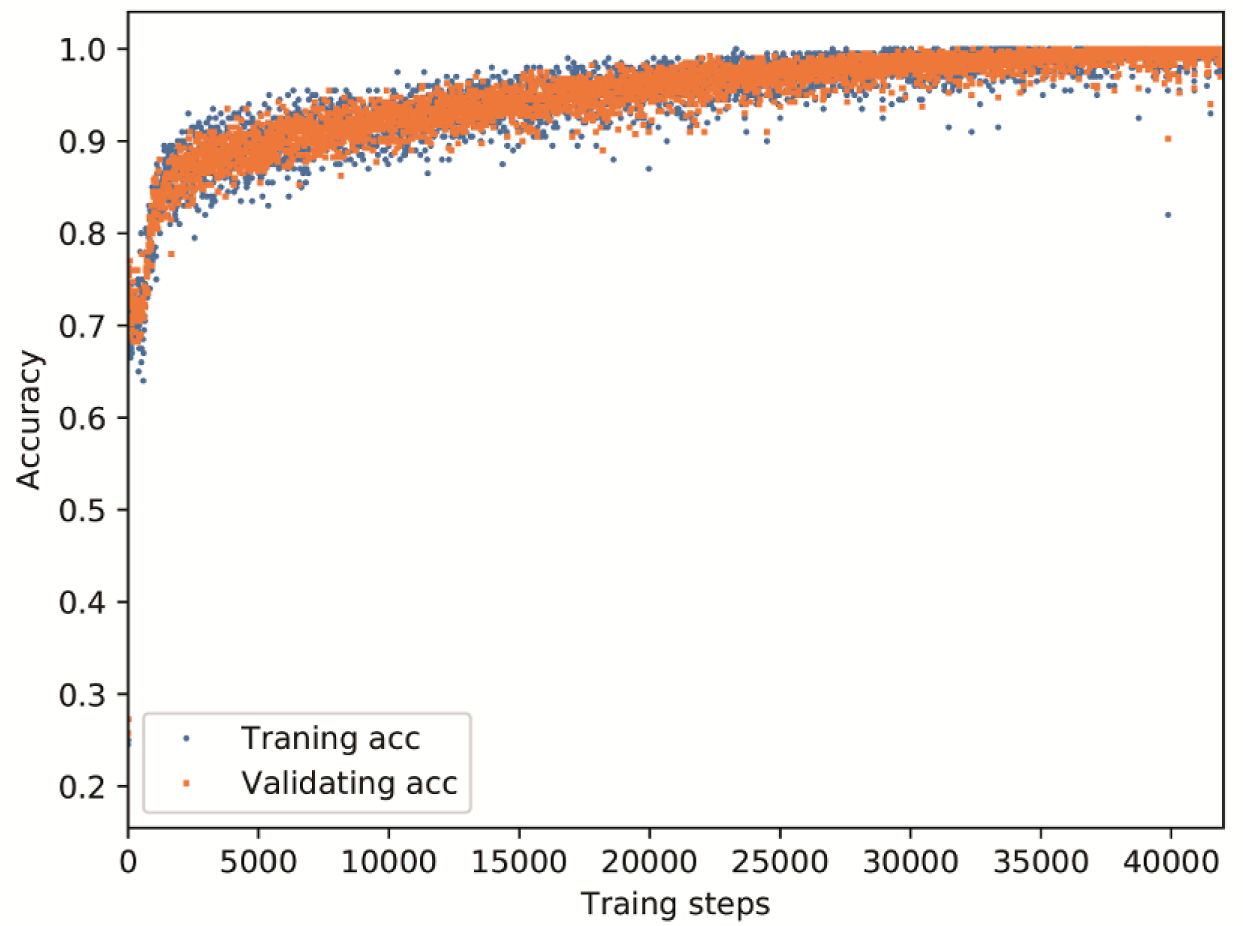
Training process of Deep-RBPPred. The training set is random divided into three parts, 200 training sequences, 600 validating sequences and other, in each training step. The model was trained with 200 training sequences and validated with 600 sequences. The training process is stopped at 42,000 steps for the training ACC and validating ACC are closed to 1 and waved as the training steps.

### Model selection

For the deep learning, an important problem is to select an appropriate model. Figure 2 shows hundreds of models may be used as the final model. In order to select a best model to predict the RBP, we test all the models in the testing dataset. We filter the models with the average ACC tested in three proteomics is greater than 0.88, training ACC and validating ACC are both greater than 0.95. For 42,000 models, only 72 models are kept. The ACC of 72 models tested in three proteomics are shown in Figure 3. The best average ACC of all the models is 0.8892, which is chosen as the final model for the following study. In Figure 3, it also shows the all the models performances best in the H. sapiens, then in the A. thaliana and the worst in the S. cerevisiae. This performances difference is also found in SVM model of RBPPred (Zhang and Liu 2017). This phenomenon may be caused by the training set which implies new sequence should be added to train the model.

**Figure 3.**
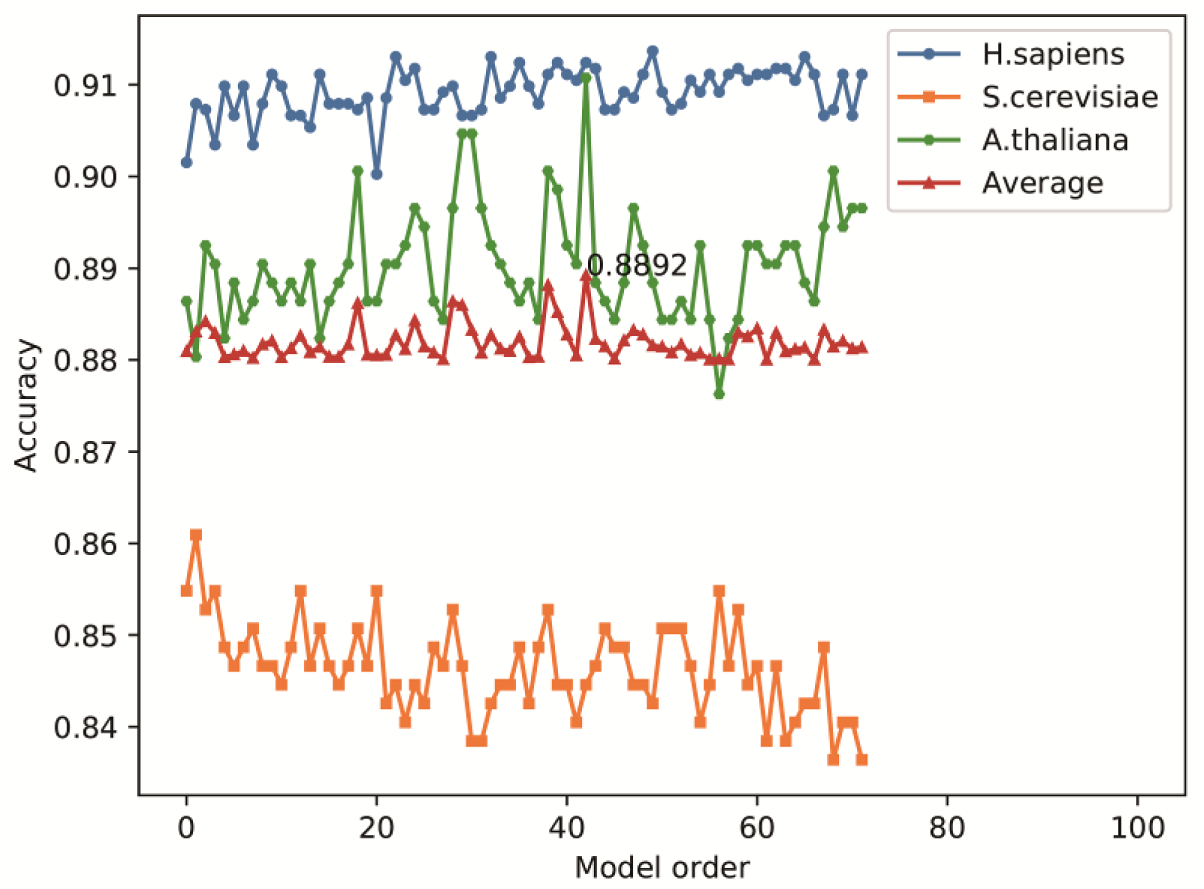
Model selection in training process. Model order is plotted against accuracy in H. sapiens, S. cerevisiae, A. thaliana proteomics of testing set and average. The best average of H.sapiens, S.cerevisiae and A.thaliana accuracy is 0.8892.

### Performance in testing data and comparison with RBPPred

For better understanding the performance of final model in the testing dataset, we evaluate the model with MCC, F-measure and ACC. We also make a comparison with RBPPred. Figure S1 shows the distribution of probability score testing in H. sapiens, S. cerevisiae and A. thaliana proteomics in testing set. Almost protein probability scores are distributed in the two ends of the figure. This distribution implies that 0.5 is an appropriate probability score to identity the RBP or non-RBP. Figure S2 describes the relationship between F-measure (MCC, ACC) and probability score. The values of MCC (F-measure) are 0.8141 (0.9034), 0.6077 (0.7853) and 0.573 (0.9058) respectively when the score cutoff is set to 0.5. The MCC tested in H. sapiens and A. thaliana are better than RBPPred (Zhang and Liu 2017). Figure 4 shows the ROC curve of Deep-RBPPred tested in three proteomics. These results indicate that Deep-RBPPred is better than RBPPred.

**Figure 4.**
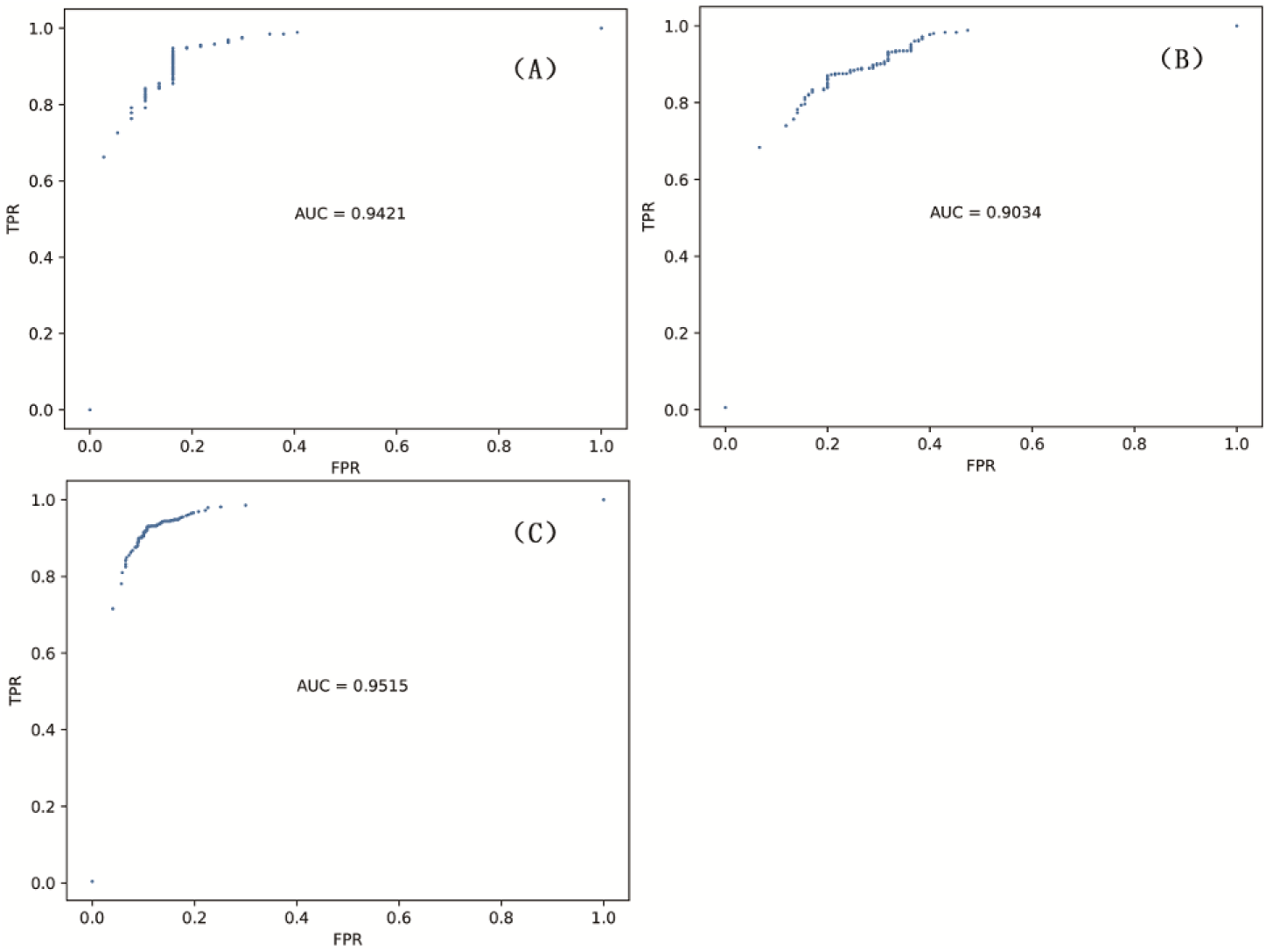
ROC of model testing in testing dataset. The AUC are 0.9421 (A), 0.9034 (B) and 0.9515 (C) for A. thaliana, S. cerevisiae and and H. sapiens respectively.

### Application to Gerstberger-1538

Recently, 1452 RNA binding proteins in human have been identified from Pfam (Finn et al. 2016). We used Gerstberger-1538 from RBPPred and verified our model in this dataset. Figure 5 shows the distribution of probability score. The SN is 0.9038 which is about 18% higher than RBPPred. This result indicates that Deep-RBPPred has a good generalization ability.

**Figure 5.**
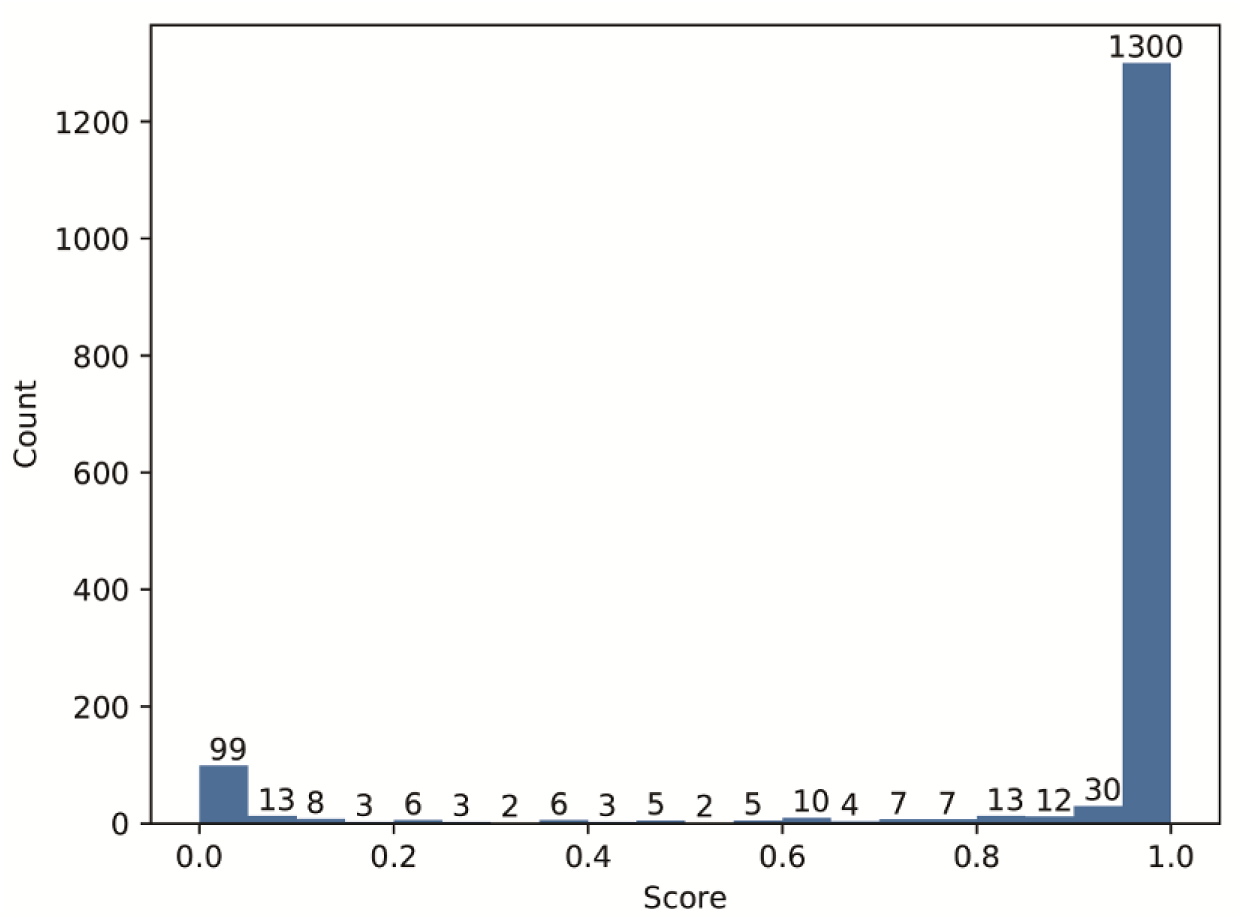
Distribution of probability score testing in Gerstberger-1538.

### Computational time

Running time is an important metric to measure a model. We list the computational time of Deep-RBPPred in Table 1. The table shows that Deep-RBPPred is a very fast RBP prediction model. Here we do not list the computational time of RBPPred because it cost much computational time. Comparing to RBPPred, Deep-RBPPred predicts RBP without using blast to generate PSSM matrix which costs much computational time. Take advantage of less computational time, Deep-RBPPred is used to studying in proteome scale.

**Table 1.**
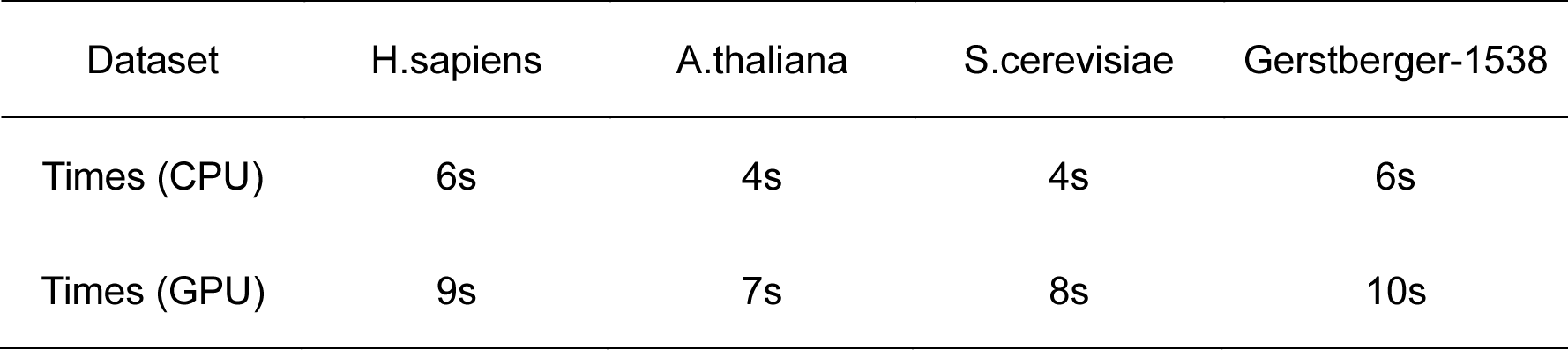
Computational time of Deep-RBPPred running in Centos with Intel(R) Xeon(R) CPU E5-2620 v2 @ 2.10GHz or GeForce GTX 980.

| Dataset | H.sapiens | A.thaliana | S.cerevisiae | Gerstberger-1538 |
| --- | --- | --- | --- | --- |
| Times (CPU) | 6s | 4s | 4s | 6s |
| Times (GPU) | 9s | 7s | 8s | 10s |

### Capacity of predicting new RBPs

To test the predicting ability of Deep-RBPPred on new RBPs, we collected 299 new RBPs created between 2016-05-24 (consistent with RBPPred) and 2017-09-27 in Uniprot. Then Deep-RBPPred and RBPPred are both run on the dataset. 94.65% of 299 new RBPs are correctly predicted by Deep-RBPPred, but only 86.87% are predicted by RBPPred. Deep-RBPPred performs about 8% better than RBPPred. One protein (Uniprot id: P0DOC6) cannot be calculated by RBPPred for that no protein sequences can be found by Blast (Altschul et al. 1990). These results indicate that Deep-RBPPRed has a better performance and more widely application than RBPPRed.

### RBPs prediction in the 30 proteomics

One of the characteristic of our approach is running fast. Testing in 1538 protein sequences only costs 10s when is run in GTX 980. Hence, Deep-RBPPred is used to widely test in proteome scale. Figure S3 shows the predicting result of deep-RBPPred and RBPPred in H.sapiens, S.cerevisiae and A.thaliana proteomics derived from the Uniprot (reviewed). Deep-RBPPred predicts 16648, 5049 and 11551 RBPs in H.sapiens, S. cerevisiae and A. thaliana proteomics respectively. RBPPred predicts 13027, 3850 and 8277 RBPs in these proteomics. 4369, 1543 and 4089 RBPs are both predicted by Deep-RBPPred and RBPPred. In all, Deep-RBPPred predicts more RBPs than RBPPred, which is unanimous to the result in Gerstberger-1538. For more widely predicting our model, we collected 30 whole proteomics including both reviewed and unreviewed proteins downloaded from Uniprot website. Figure 6 shows the predicting result of Deep-RBPPred. We found an interesting result that the rate of RBPs in eukaryotes proteome is higher than bacteria. This phenomenon indicates RBPs in eukaryotes are successful than bacteria in terms of numbers of non-RBPs. This result implies RBPs may function in more complex cellular process. For human, mouse, A.thaliana, rat, zebrafish and E.coli (strain K12) proteome, we also calculated the SN. The results are show in Table 2. For human, the whole proteome includes 2753 RBPs. Deep-RBPPred obtains SN 0.8373. So, we can estimate the total number of RBPs in human is about 34047(71591*0.568*0.8373). This number is higher than any experiment discovered RBPs in human. The same analysis goes for other organisms. The above results indicate the total number of RBPs may be more than we discovered at present.

**Figure 6.**
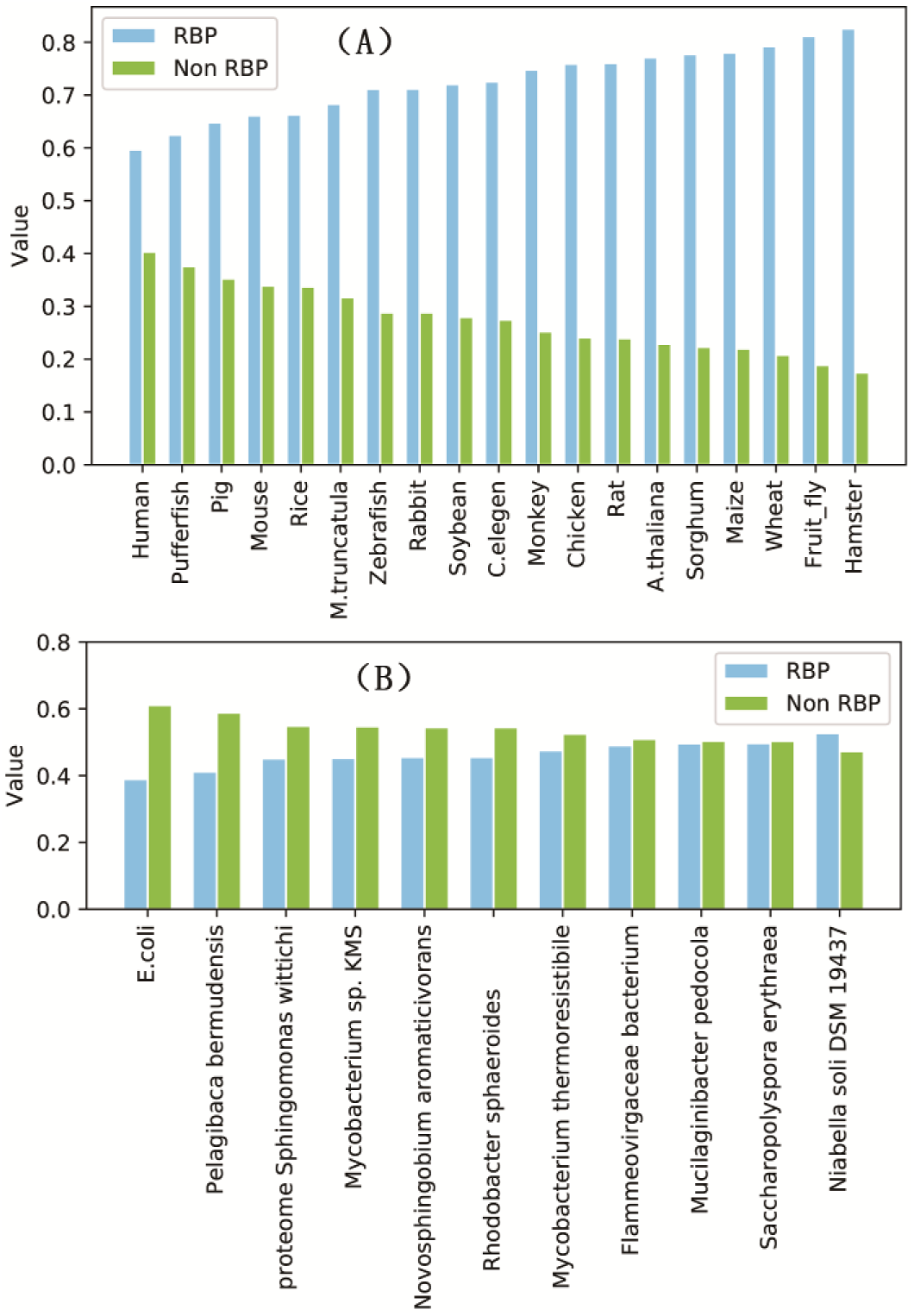
RBP prediction of Deep-RBPPred in 19 eukaryotes proteomics (A) and 11 bacteria proteomics (B). *Value is defined as the number of RBP (non-RBP)/ the number of all proteins in a proteome*.

**Table 2.**
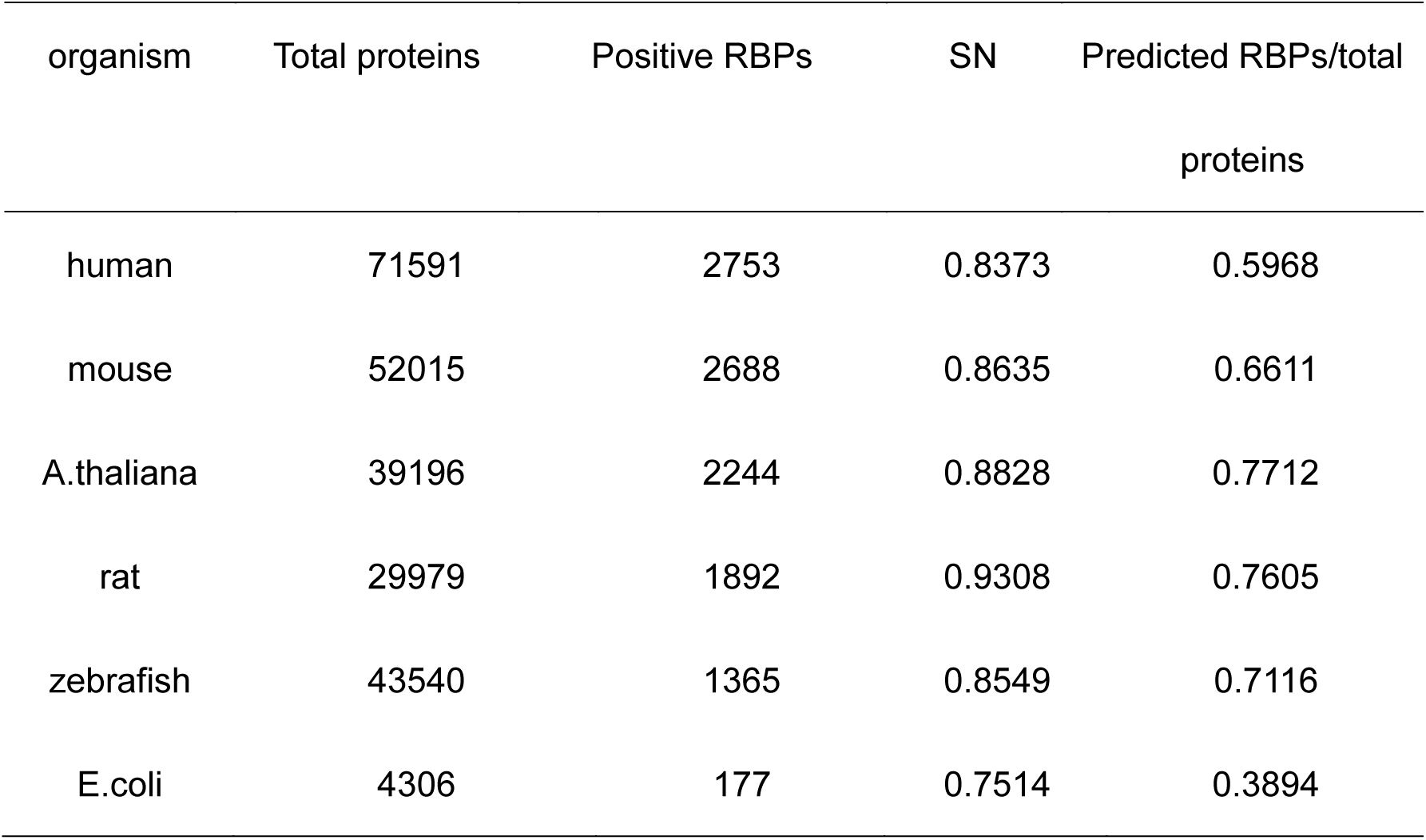
RBP prediction in 6 proteomics.

| organism | Total proteins | Positive RBPs | SN | Predicted RBPs/total<br>proteins |
| --- | --- | --- | --- | --- |
| human | 71591 | 2753 | 0.8373 | 0.5968 |
| mouse | 52015 | 2688 | 0.8635 | 0.6611 |
| A.thaliana | 39196 | 2244 | 0.8828 | 0.7712 |
| rat | 29979 | 1892 | 0.9308 | 0.7605 |
| zebrafish | 43540 | 1365 | 0.8549 | 0.7116 |
| E.coli | 4306 | 177 | 0.7514 | 0.3894 |

## Discussion

In this study, we develop an RBP predicting model based on CNN which only needs hydrophobicity, normalized van der waals volume, polarity and polarizability, solvent accessibility, charge and polarity of side chain of protein sequence. The result tested on H. sapiens, S. cerevisiae, A. thaliana proteomics shows that our model performances as good as RBPPred which is the best accuracy model so far. We also verify our model in Gerstberger-1538 which is extracted from Pfam containing experimental validated and computational predicted. In the testing in Gerstberger-1538, the SN of our model is 18% higher than RBPPred. Predicting in 3 protein proteomics derived from Uniprot (reviewed), Deep-RBPPred predicts more RBPs than RBPPred, and almost RBPs predicted by RBPPred are also predicted by Deep-RBPPred. The SN tested on 299 new RBPS is about 8% higher than RBPPred. Finally, we predicted the RBPs in 30 whole reference proteomics (both reviewed and unreviewed). We found an interesting result that the rate of RBPs in eukaryotes proteome is higher than bacteria. This result is needed to widely verify. The results also indicate the total number of RBPs may be more than we discovered at present.

### Software availability

Deep-RBPPred is written in the python, availability as an open source tool at http://www.rnabinding.com/Deep_RBPPred/Deep-RBPPred.html.

## ACKNOWLEDGMENTS

We thank the National Supercomputer Center in Guangzhou for support of computing resources. This work has been supported by the Fundamental Research Funds for the Central Universities [2016YXMS017] and the Special Program for Applied Research on Super Computation of the NSFC-Guangdong Joint Fund (the second phase) under Grant No.U1501501.

---

**Figure S1.**
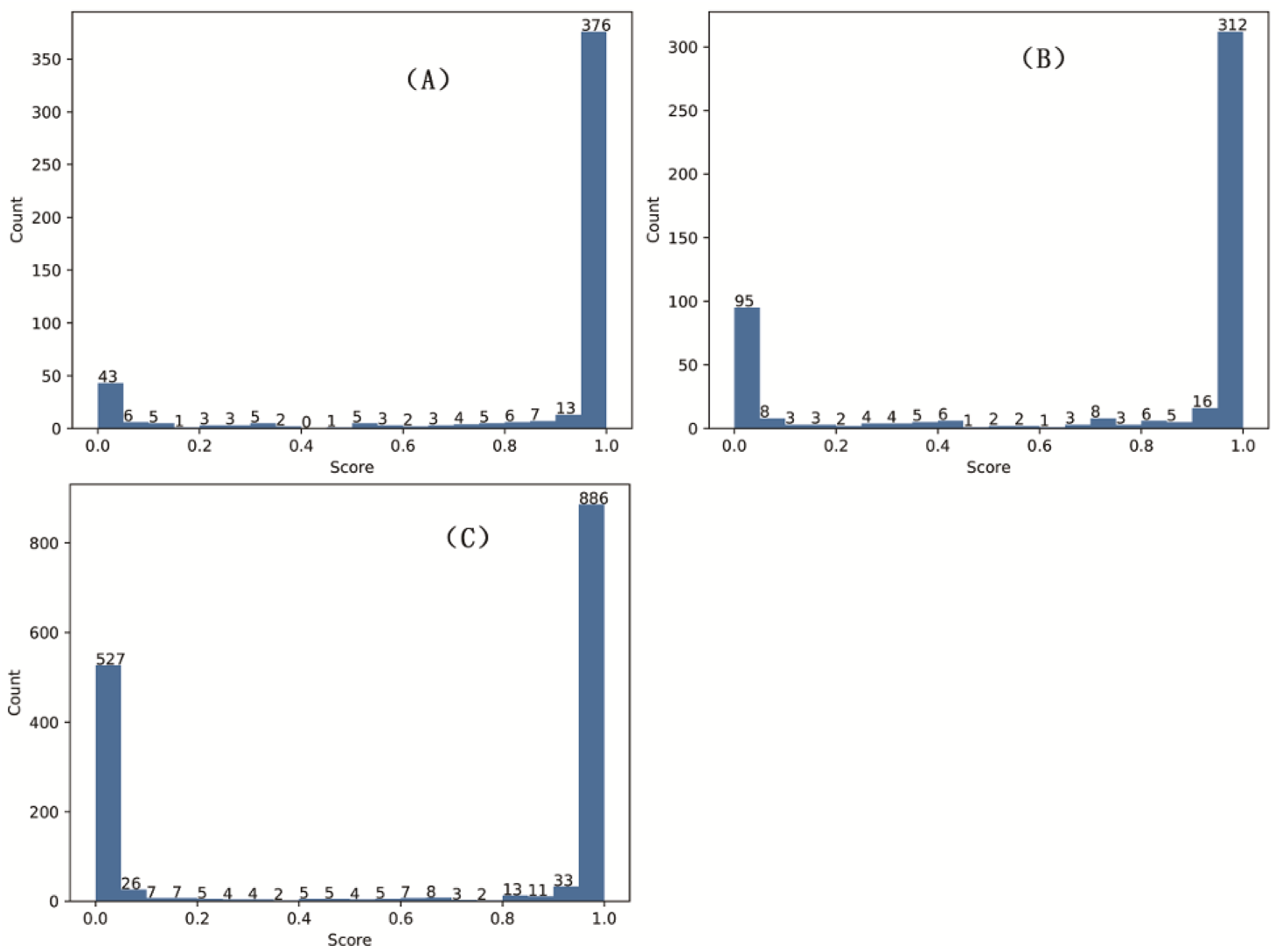
Distribution of probability score testing in in A. thaliana (A), S. cerevisiae (B) and H. sapiens (C) proteomics of testing set.

**Figure S2.**
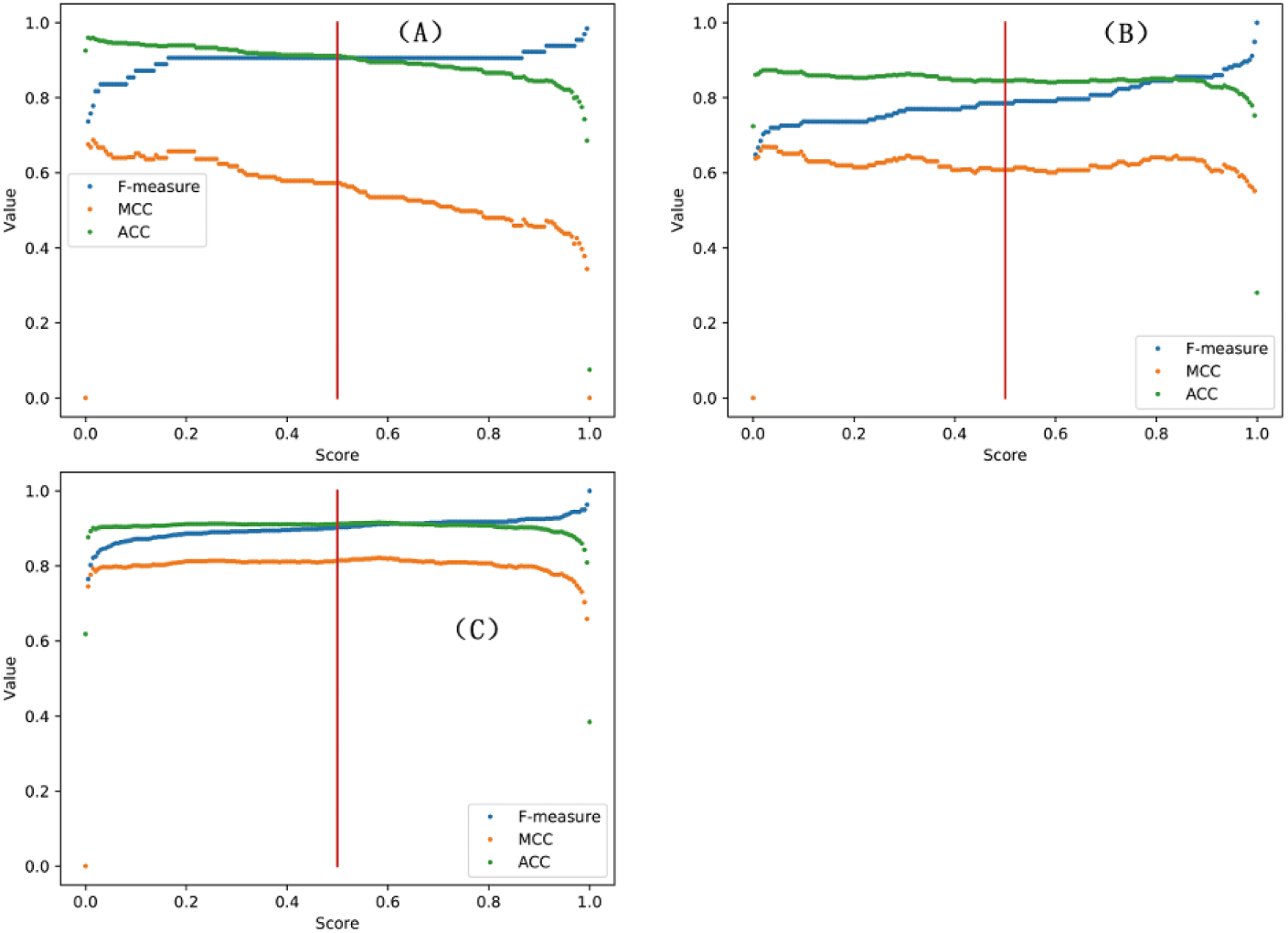
F-meaure, MCC and ACC vs probability score. F-meaure, MCC and ACC are plotted against probability score for A. thaliana (A), S. cerevisiae (B) and H. sapiens (C) proteomics of testing set. The red vertical line indicates the status where the probability score is chosen as 0.5 to identify the RBP.

**Figure S3.**
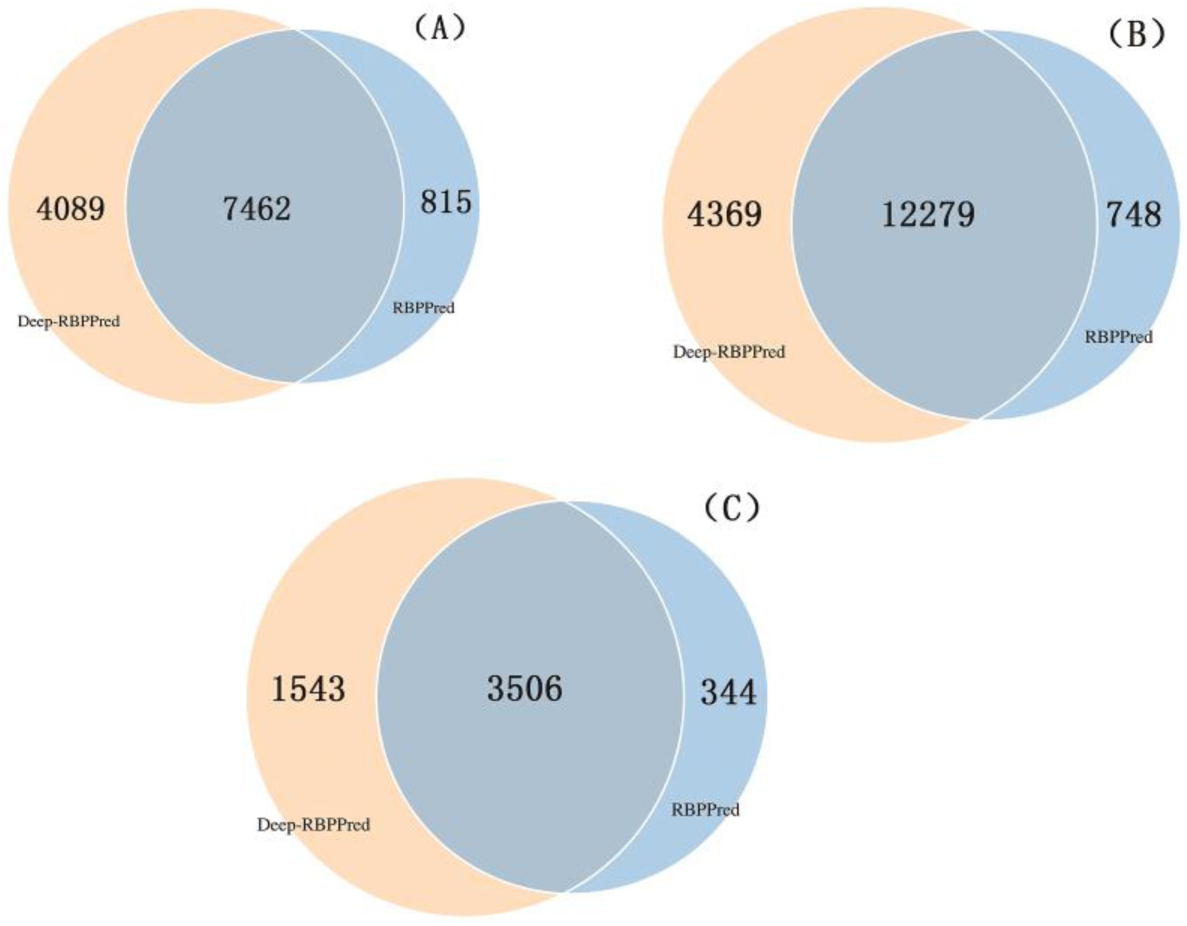
Comparison of RBPPred and deep-RBPPred predicting in A.thaliana (A), H.sapiens (B) and S.cerevisiae (C) proteomics.

